# SARS-CoV-2 infection unevenly impacts metabolism in the coronal periphery of the lungs

**DOI:** 10.1101/2024.05.22.595414

**Authors:** Jarrod Laro, Biyun Xue, Jian Zheng, Monica Ness, Stanley Perlman, Laura-Isobel McCall

## Abstract

**Highlights:**

- COVID-19 significantly decreases amino acids, fatty acids, and most eicosanoids
- SARS-CoV-2 preferentially localizes to central lung tissue
- Metabolic disturbance is highest in peripheral tissue, not central like viral load
- Spatial metabolomics allows detection of metabolites not altered overall

SARS-CoV-2, the virus responsible for COVID-19, is a highly contagious virus that can lead to hospitalization and death. COVID-19 is characterized by its involvement in the lungs, particularly the lower lobes. To improve patient outcomes and treatment options, a better understanding of how SARS-CoV-2 impacts the body, particularly the lower respiratory system, is required. In this study, we sought to understand the spatial impact of COVID-19 on the lungs of mice infected with mouse-adapted SARS2-N501Y_MA30_. Overall, infection caused a decrease in fatty acids, amino acids, and most eicosanoids. When analyzed by segment, viral loads were highest in central lung tissue, while metabolic disturbance was highest in peripheral tissue. Infected peripheral lung tissue was characterized by lower levels of fatty acids and amino acids when compared to central lung tissue. This study highlights the spatial impacts of SARS-CoV-2 and helps explain why peripheral lung tissue is most damaged by COVID-19.

## Introduction

Severe acute respiratory syndrome coronavirus 2 (SARS-CoV-2) is a respiratory virus that causes coronavirus disease 2019 (COVID-19). COVID-19 can have a wide range of severity: some patients remain asymptomatic while others experience mild symptoms such as fatigue, nausea, and a loss of taste and smell. In dire cases, patients experience hypoxia and acute respiratory disease syndrome (ARDS), which can ultimately lead to death. Indeed, by the end of 2023, the World Health Organization recorded over 700 million cases of COVID-19 worldwide, with 7 million deaths being attributed directly to the infection or its complications.^1^ SARS-CoV-2 is highly contagious and mutates rapidly, explaining its continued prevalence and appearance of new variants despite widespread availability of vaccines.^2^ Even after acute SARS-CoV-2 infection resolves, a substantial fraction of patients report continued feelings of fatigue, breathlessness, chest pains, and the onset of conditions such as type 2 diabetes and postural orthostatic tachycardia syndrome (POTS).^3,4^ These post-acute sequelae, most commonly referred to as “long COVID”, are poorly characterized, with little understanding of why they occur, how to effectively diagnosis them, or how to prevent them.^5^ Due to its continued presence, it is crucial to understand how SARS-CoV-2 impacts the host, particularly the lower respiratory system, to improve patient outcomes and to develop better treatments and prophylactics.

The metabolome, the collection of molecules with a molecular weight under 1500 Daltons present in an organism, is extremely sensitive to small changes caused by age, gender, sex, diet, illness, etc., making analysis of the metabolome a useful tool for understanding the impact and mechanism of disease.^6^ Typically, metabolomic studies investigate a biofluid, a small sampling of an organ, or the organ as whole. The study of the serum metabolome has been helpful in discerning some of the impact of conditions such as Chagas Disease, influenza, and COVID-19, but the study of biofluids is unable to capture the heterogeneity present within biofluid or organs.^7,8^ Indeed, metabolomic studies that have segmented the heart, lungs, liver, et cetera, have identified differences in metabolism between different parts of the same organ.^9–12^ While SARS-CoV-2 primarily targets the lungs, the specific affected lung segments can vary based on the patient. Tissue damage can present in a single lobe of the lung or all lobes, be peripherally or centrally distributed in the transversal plane, and cause ground glass opacities or consolidations.^13–16^ Despite the heterogeneous presentation of the condition, the lower segments of the coronal plane are the ones most commonly impacted by SARS-CoV-2 in humans, with the majority of cases showing abnormalities in the periphery of the transverse plane of these segments.^14,16–18^ While many studies have examined the impact of COVID-19 on the metabolome (e.g. ^19–22)^, none have attempted to explain the localized impact of SARS-CoV-2 through the use of spatial metabolomics. Additionally, due to its ease of collection in clinical settings, the majority of COVID-19 studies focus on common biofluids like plasma and urine (for example, ^23–27)^. While plasma provides information on the biological system as a whole, it cannot capture localized, distinct effects of a condition within individual organs.

To address these gaps, we investigated the impact of COVID-19 by performing a systematic spatial metabolomic analysis of lung tissues from mice infected with the mouse-adapted SARS2-N501Y_MA30_ strain^28^ at a sublethal or lethal dose (1000 plaque forming units and 5000 plaque forming units, respectively) to mimic different levels of COVID-19 severity five days post-infection (DPI). To identify potential markers of post-acute sequelae, we analyzed lung tissues twenty days post-infection from mice infected with 1000 plaque forming units. Through this approach, we identified localized effects of COVID-19 on lung metabolism and preferential localization of SARS-CoV-2 in lung tissue.

## Results

### SARS-CoV-2 infection causes substantial metabolic changes that persist at least 20 days after infection

To examine the impact of COVID-19 on lung metabolism, five experimental groups were used: mock-infected mice euthanized 5 DPI (Mock5) or 20 DPI (Mock20), mice infected with 1000 or 5000 plaque-forming units of SARS2-N501Y_MA30_ and euthanized 5 DPI (D5-1000 and D5-5000), and mice infected with 1000 plaque-forming units of SARS2-N501Y_MA30_ and euthanized 20 DPI (D20-1000). Initially, the overall metabolic differences between each disease state were analyzed irrespective of lung position. We observed that the mock-infected mice possessed strong metabolic differences from infected mice (figure 1A; Pseudo-F > 60 for D5-1000, D5-5000, and D20-2000 when compared to the time-matched mock-infected control). D5-5000 demonstrated the largest metabolic perturbation relative to their time-matched mock-infected control, followed by D5-1000 and D20-1000 (Supplemental Figure 1). D5-1000 and D5-5000 mice also displayed strong metabolic differences from each other (Pseudo-F = 23.1, q-value < 0.001), made more apparent when analyzing only those two groups (Figure 1B). While D20-1000 mice had clear metabolic differences from mock-infected mice, they displayed a wide range of phenotypes, with some individual mice clustering closer to uninfected mice while others clustered with the D5-1000 and D5-5000 mice (Figure 1A; Supplemental Figure 2A). Metabolic differences between mice were not clearly associated with viral load measured from lung tissue (Supplemental Figure 3).

**Figure 1.**
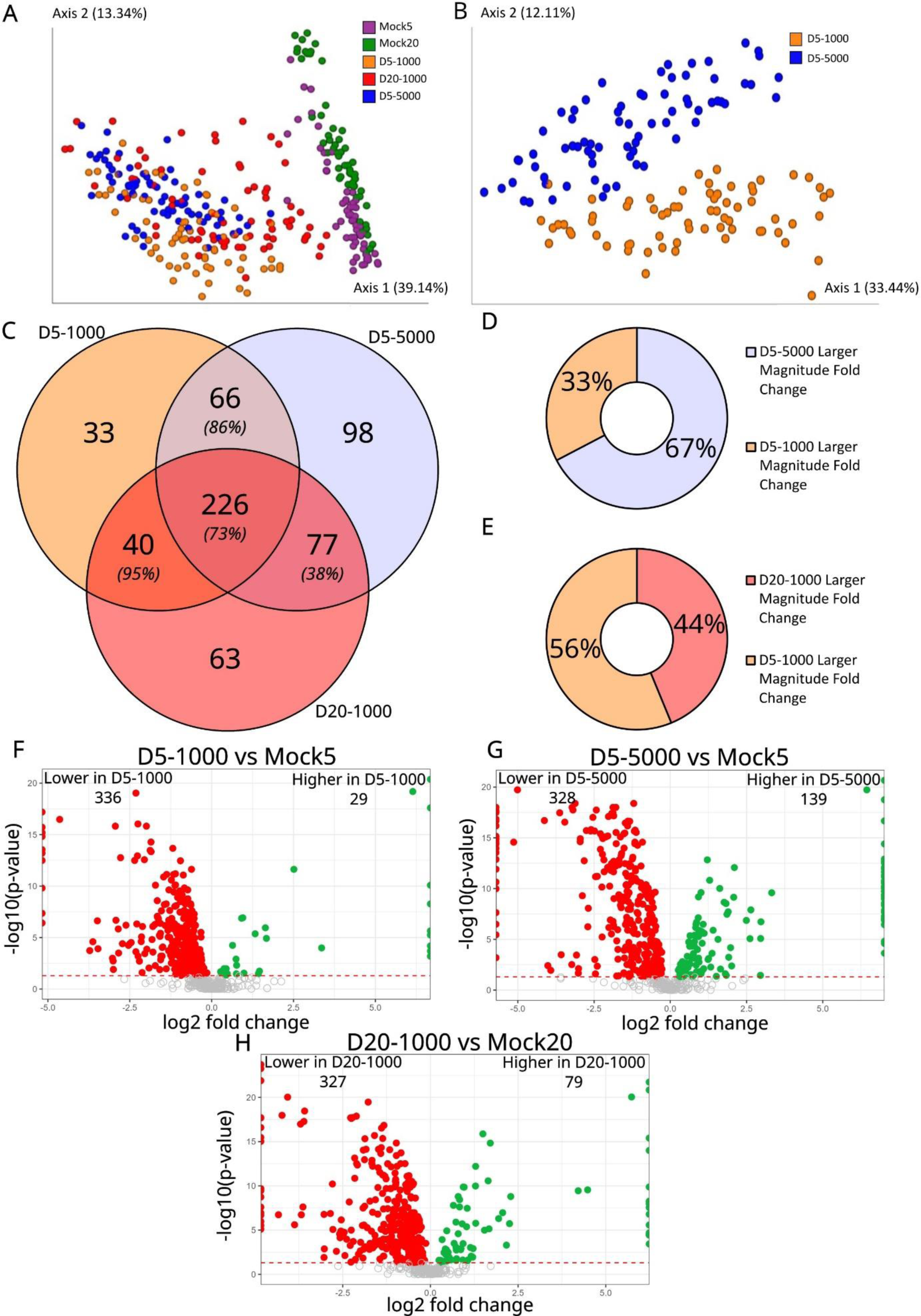
SARS-CoV-2 infected mice show strong metabolic differences from mock-infected mice overall in the lung. (A) Principal coordinate analysis for all infection states. (B) Principal coordinate analysis for D5-1000 vs D5-5000. (C) Overlap of the metabolites significantly impacted in each of the infected mice when compared to timepoint-matched mock-infected controls. Italicized percentages in each overlap are the percent of metabolites that all have the same direction of change. (D) Comparison of the magnitude of log2 fold changes between annotated, significant metabolites in both D5-5000 and D5-1000 mice that have the same direction of fold change (n = 254). (E) Comparison of the magnitude of log2 fold change between annotated, significant metabolites in both D20-1000 and D5-1000 mice that have the same direction of fold change (n = 217). (F-H) Volcano plots of annotated metabolites for (F) D5-1000, (G) D5-5000, (H) and D20-1000 groups compared to timepoint-matched mock-infected controls. Significant metabolites (FDR-adjusted p-value < 0.05) are filled-in, colored data points. Red dots, fold change<1 and green dots, fold change> 1. Gray, unfilled data points are metabolite features that were not significantly different.

### SARS-CoV-2 infection decreases major molecular families in the lung

To identify individual metabolites impacted by COVID-19, p-values were calculated by comparing each infected group with their time-matched mock-infected control and then false-discovery rate (FDR) adjusted; fold change values for each disease state were calculated by dividing the median total ion current (TIC) normalized signal intensity of the metabolite in the infected group by the median TIC-normalized signal intensity of the metabolite in the time-matched mock-infected controls. The majority of significantly altered metabolites (FDR-adjusted p-value < 0.05) were shared between all three disease states (Figure 1C). Overall, metabolites affected under all three infection conditions, shared between D5-1000 and D20-1000, or shared between D5-1000 and D5-5000, predominantly showed the same direction of change, highlighting reproducibility of the metabolic impact of infection. Fold change differences to the time-matched mock-infected control were more pronounced for two-thirds of the metabolites significantly impacted in D5-5000 mice when compared to D5-1000 mice (Figure 1D; χ^2^ = 30.488, p-value < 0.00001). No significant difference was observed between the magnitude of the fold change differences in D5-1000 mice versus D20-1000 mice (Figure 1E; χ^2^ = 3.613, p-value = 0.06). The majority of significantly altered metabolites exhibited a decrease relative to the mock control, with 92%, 70%, and 81% of metabolites showing a significant decrease in D5-1000, D5-5000, and D20-1000 mice respectively, compared to time-matched mock-infected controls (Figure 1F-H).

Overall, infected lung tissue across all states had significantly altered levels of amino acids, fatty acids, eicosanoids, acylcarnitines, and purines, amongst other molecular families (Figure 2A-C). Amino acids were decreased across all disease states (Figure 2A-D). D5-1000 mice exhibited significantly lower levels of glutamate, methionine, and arginine compared to mock-infected controls; in addition to those amino acids, D5-5000 mice had significantly lower levels of threonine, tyrosine, phenylalanine, and tryptophan compared to mock-infected controls. D20-1000 mice had a significant decrease in threonine, glutamate, tyrosine, and arginine. Alongside the decrease in tryptophan levels observed in D5-5000 mice, there was a significant increase in the levels of kynurenine, a well-characterized sign of inflammation due to activation of the TDO/IDO-catalyzed transformation of tryptophan into kynurenine. Kynurenine was not significantly increased in either D5-1000 or D20-1000 mice, implying lower levels of inflammation and disease severity in these groups (Supplemental Figure 4).

**Figure 2.**
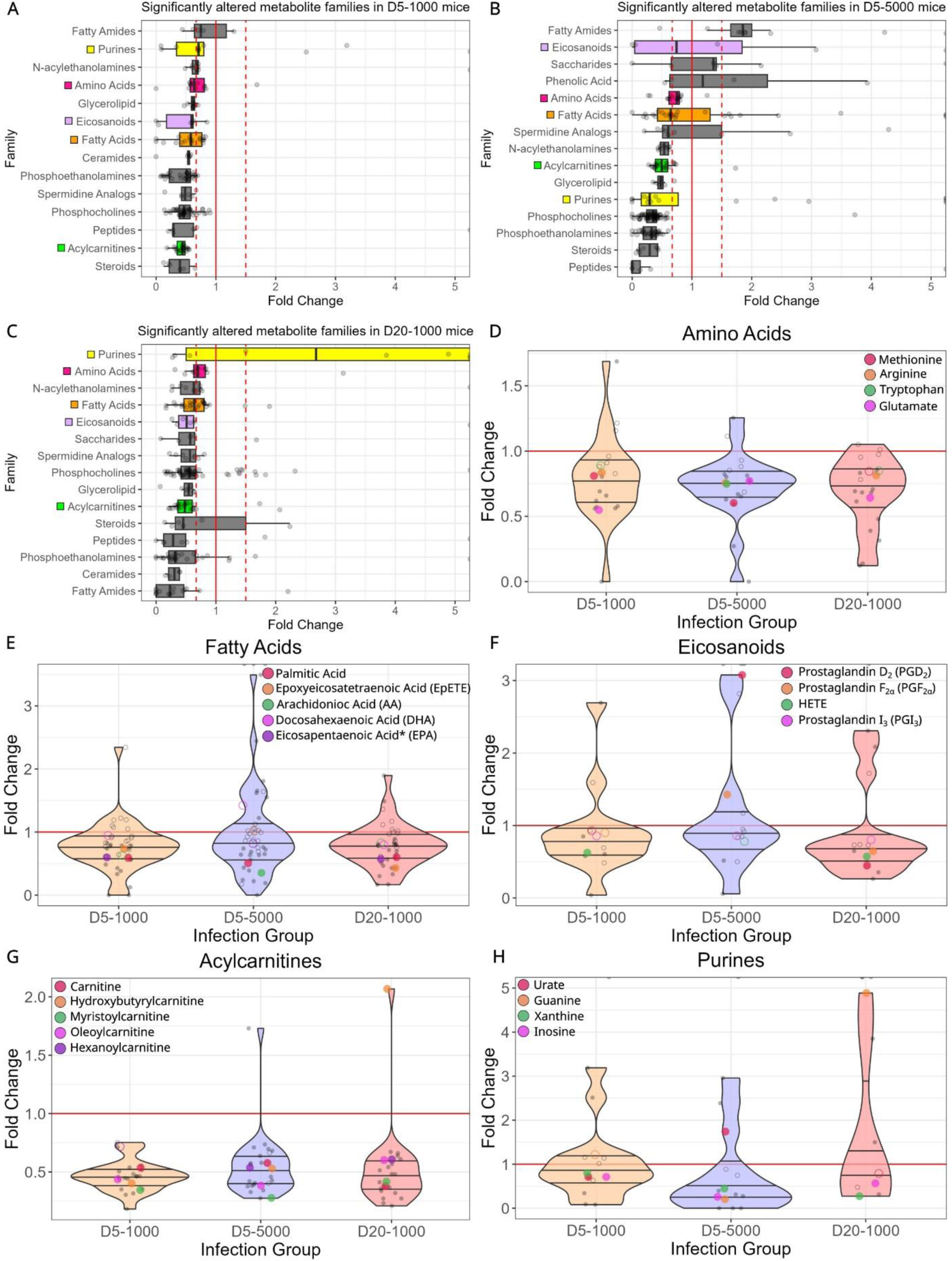
SARS-CoV-2 infection decreases major molecular families in the lung. All data represented as median fold change (A-C) Significantly altered molecular families for (A) D5-1000, (B) D5-5000, and (C) D20-1000 mice compared to timepoint-matched mock-infected controls. All families depicted had at least 4 annotated metabolites belonging to that family. Dots on the right edge of each panel represent metabolites that had fold changes greater than 5. Dashed red lines represent a fold change of 0.67 or 1.5 while the solid red line represents a fold change of 1. Families visualized in panels D-H were colored to make them easier to observe between panels A-C. (D-H) Violin plots of specific molecular families across all 3 disease states compared to timepoint matched mock-infected controls. Solid dots represent members of the family with a FDR-adjusted p-value < 0.05 while hollow dots represent members of the family with a FDR-adjusted p-value > 0.05. Red line, fold change of 1.

Fatty acid metabolism was significantly impacted in all disease states (Figure 2A-C, 2E). Palmitic acid (PA), epoxyeicosatetraenoic acid (EpETE), and arachidonic acid (AA) were all lower in D5-1000 mice and D20-1000 mice compared to mock-infected controls. *m/z* 303.23 RT 3.02 min, annotated as eicosapentaenoic acid (EPA) or an isomer (Supplemental Figure 5) was also lower in D5-1000 mice and D20-1000 mice compared to mock-infected controls. PA and AA were both significantly lower in D5-5000 mice, whereas EpETE and EPA were not significantly altered. Docosahexaenoic acid (DHA), a 22-carbon polyunsaturated ω-3 fatty acid with anti-inflammatory effects ^29–31^, was not significantly altered in any disease state. 20-carbon polyunsaturated fatty acids like AA and EPA serve as the precursor for eicosanoids, which play a key role in inflammatory responses.^32–35^ Eicosanoid products of the ω-3 fatty acid EPA, such as prostaglandin I_3_ (PGI_3_), prostaglandin E_3_, resolvins, and protectins, have anti-inflammatory effects that temper the effects of AA-derived eicosanoids.^31,36–39^ Eicosanoids were generally decreased in D5-1000 and D20-1000 mice compared to timepoint-matched mock-infected controls, but had larger fold changes in D5-5000 (Figure 2F). Multiple eicosanoids were only detected in D5-5000 tissue. Prostaglandin D_2_ (PGD_2_) and prostaglandin F2_α_ (PGF_2α_) were not significantly altered in D5-1000 mice but were significantly increased in D5-5000 mice and significantly decreased in D20-1000 mice compared to mock-infected controls. PGI_3_ was detected but not significantly altered in overall lung tissue. Indeed, PGD_2_ and PGF_2α_ had a significant, positive correlation with post-infection viral load while PGI_3_ lacked any significant correlation (Supplemental Table 1).

Acylcarnitines and their building block, carnitine, were almost ubiquitously lower across all three disease states (Figure 2A-C, 2G). Significantly altered acylcarnitines include hexanoylcarnitine (C6, medium-chain), myristoylcarnitine (C14, long chain), and oleoylcarnitine (C18:1, unsaturated long chain). Hydroxybutyrylcarnitine (C4, hydroxy short chain) was the only detected acylcarnitine with elevated levels at D20. The impact of SARS-CoV-2 infection did not appear to be acyl chain length specific. Acylcarnitines are crucial molecules for the transport of fatty acids across mitochondrial walls. Upon being transported into the mitochondria, fatty acids are able to be catabolized to produce energy. Decreased levels of acylcarnitines in tandem with lower levels of several fatty acids could be a product of increased β-fatty acid oxidation for the production of energy for viral replication.

A significant impact on purine metabolism was observed in infected mice (Figure 2A-C, 2H). Urate, the end product of purine metabolism, was significantly lower in D5-1000 mice, but significantly higher in D5-5000 mice compared to controls. Xanthine, guanine, and inosine were all significantly lower in D5-5000 mice. Interestingly, in D20-1000 mice, multiple adenosine analogs (adenosine monophosphate and deoxyadenosine monophosphate) and guanine/guanosine presented with fold change increases of up to 20-fold when compared to Mock20 mice or were only detected in the D20-1000 mice. When combining D5-1000 and D5-5000 mice, purines demonstrated a negative correlation with viral loads while urate had a positive correlation (Supplemental Table 1).

### Post-infection SARS-CoV-2 viral load does not correlate with the intensity of metabolic disturbance

During initial collection of lung tissue prior to metabolite extraction and LC-MS analysis, each lung was cut into 11 lung pieces plus the trachea based on the biological segmentation of mouse lung lobes (Figure 3A/B; positions 1-5, left lung; positions 6-7, right superior lobe; position 8, right middle lobe; positions 9-10, right inferior lobe; position 11, post-caval lobe; position 12, trachea). Seeking to better understand the localization of COVID-19 effects, we categorized the 11 lung segments as either central or peripheral lung tissue. Lung segments P2-P4 and P7-P9 are from the central region of the lung, while segments P1, P5, P6, and P10 are from the peripheral region of the lung (Figure 3C). For this analysis, we did not include the post-caval mouse lung lobe (P11) due to a lack of a human analog. For all five experimental groups, there was a significant difference between the overall metabolome of the central positions and the peripheral positions, with D5-5000 mice showing the largest differences and Mock5 mice showing the smallest differences (Supplemental Table 2). As expected, D5-5000 mice had significantly higher viral loads than D5-1000 and D20-1000 mice (Figure 3F; p < 1.095e-11, Supplemental Table 3), matching the larger metabolic differences observed between D5-5000 mice and Mock5 mice. D5-1000 mice had significantly higher viral loads than D20-1000 mice (Figures 3D and 3H; p = 0.007, Supplemental Table 3). Central lung tissue had a higher median viral load than peripheral lung tissue in all three disease states. Despite this, peripheral lung tissue had significant higher levels of metabolic disturbance than central lung tissue in D5-1000 and D5-5000 (p = 0.007 and p = 0.03, respectively, Supplemental Table 3) In fact, there was no significant correlation between viral load and pseudo-F value (Supplemental Figure 6). All infected lung positions had significant differences in metabolism when compared to their time-matched mock-infected controls (Figures 3E, 3G, and 3I, Supplemental Table 3). When comparing D20-1000 mice to D5-1000 mice, upper lung segments (P1-P2, and P6-P8) did not show a statistically significant difference in metabolome and trended further away from mock-infected mice than lower lung segments, suggesting a slower recovery in the upper segments of the lungs (Supplemental Figure 2B; Supplemental Table 4). Additionally, lung segments located in the same relative positions on the left and right lungs (e.g. position P1 on the left lung and segment P6 on the right lung) exhibited remarkably similar metabolic profiles to each other in both infected and mock-infected mice (Supplemental Table 5).

**Figure 3.**
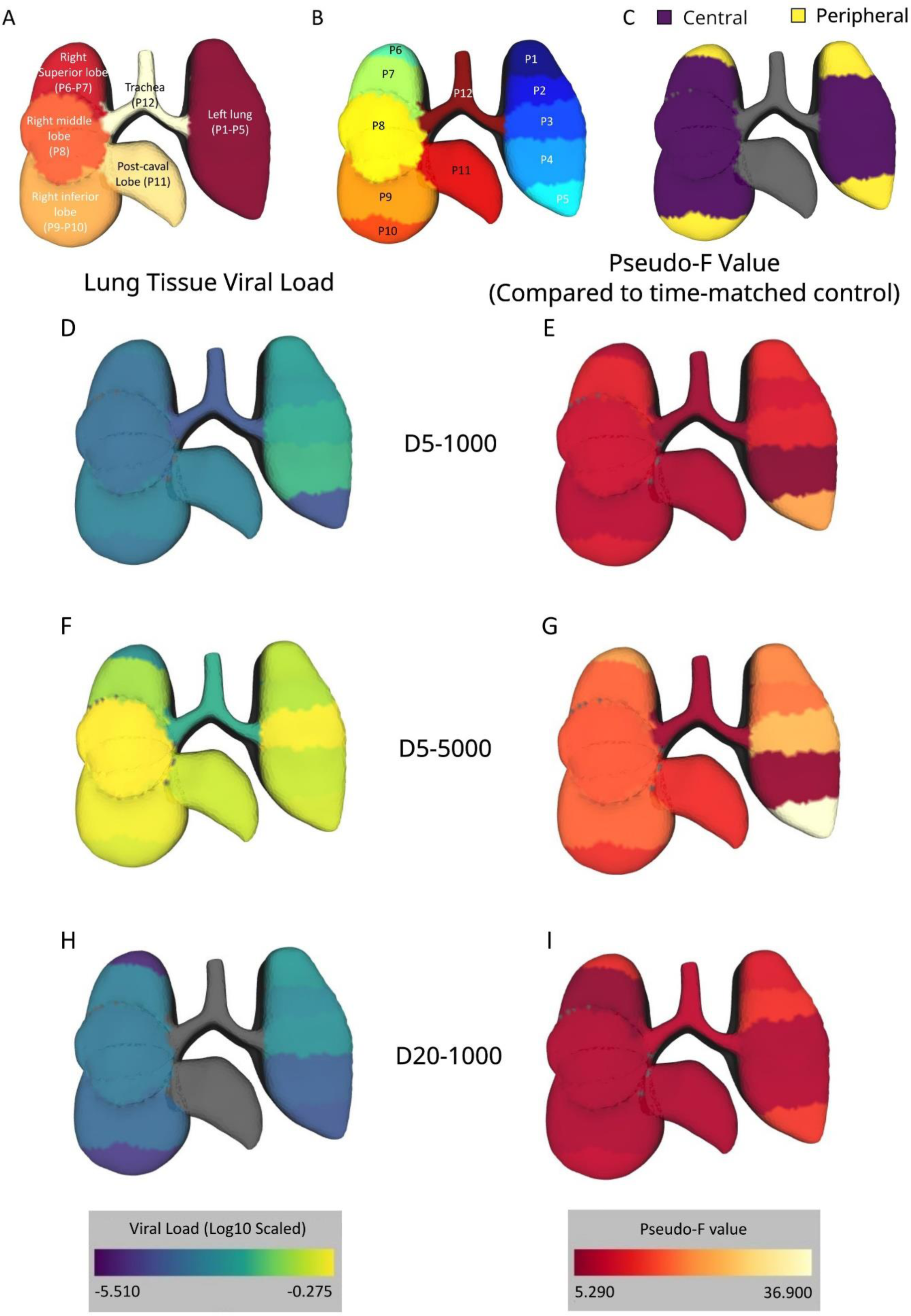
Spatial and temporal impact of SARS-CoV-2 infection on the lungs. (A) Biological segmentation of mouse lung lobes, one color per segment. (B) Experimental segmentation approach for each mouse lung. Colors show the border between the segments. (C) Classification of central (purple) vs peripheral (yellow) lung segments in the coronal plane. Position P11 and P12 are grayed out because they were not included in this analysis. (D, F, H) Log-transformed median viral loads at each position for (D) D5-1000 mice, (F) D5-5000 mice, and (H) D20-1000 mice. Gray lobes represent a median viral load of 0 (below limit of detection). Common scale. (E,G,I) PERMANOVA pseudo-F values at each position of the lung. Pseudo-F values are a measure of the overall difference between samples, with larger values representing larger differences. Each color represents the pseudo-F value at each position between the infected samples and their respective time matched mock-infected sample at that position for (E) D5-1000 mice, (G) D5-5000 mice, and (I) D20-1000 mice. Common scale. Raw data for D-I can be found in Supplemental Table 3.

### SARS-CoV-2 infection has differential localized metabolic effects on the metabolome of peripheral and central lung tissue

To investigate the impact of COVID-19 on individual metabolites, fold difference and p-value calculations were performed on the mock-normalized signal intensities for peripheral vs central lung segments. We then identified metabolites that had significantly different normalized fold differences between the peripheral and central lobes. D5-1000 mice had 111 significantly altered metabolites, D5-5000 mice had 216 significantly altered metabolites, and D20-1000 mice had 152 significantly altered metabolites (Figure 4A). 49%, 39%, and 48% of these metabolites, respectively, were only significantly altered when comparing peripheral tissue to central tissue and were not classified as significant when looking at the overall lung (Figure 4B-D). While amino acids, fatty acids, eicosanoids, and purines were all significantly decreased by infection in overall D5-1000 lung tissue (Figure 2D-F;2H), they were all significantly more decreased by infection in peripheral tissue when compared to central tissue (Figure 4E). Similarly, amino acids, fatty acids, and eicosanoids were more decreased by infection in the peripheral lung tissue of D5-5000 mice (Figure 4F), and fatty acids were more decreased in the peripheral lung tissue of D20-1000 mice (Figure 4G).

**Figure 4.**
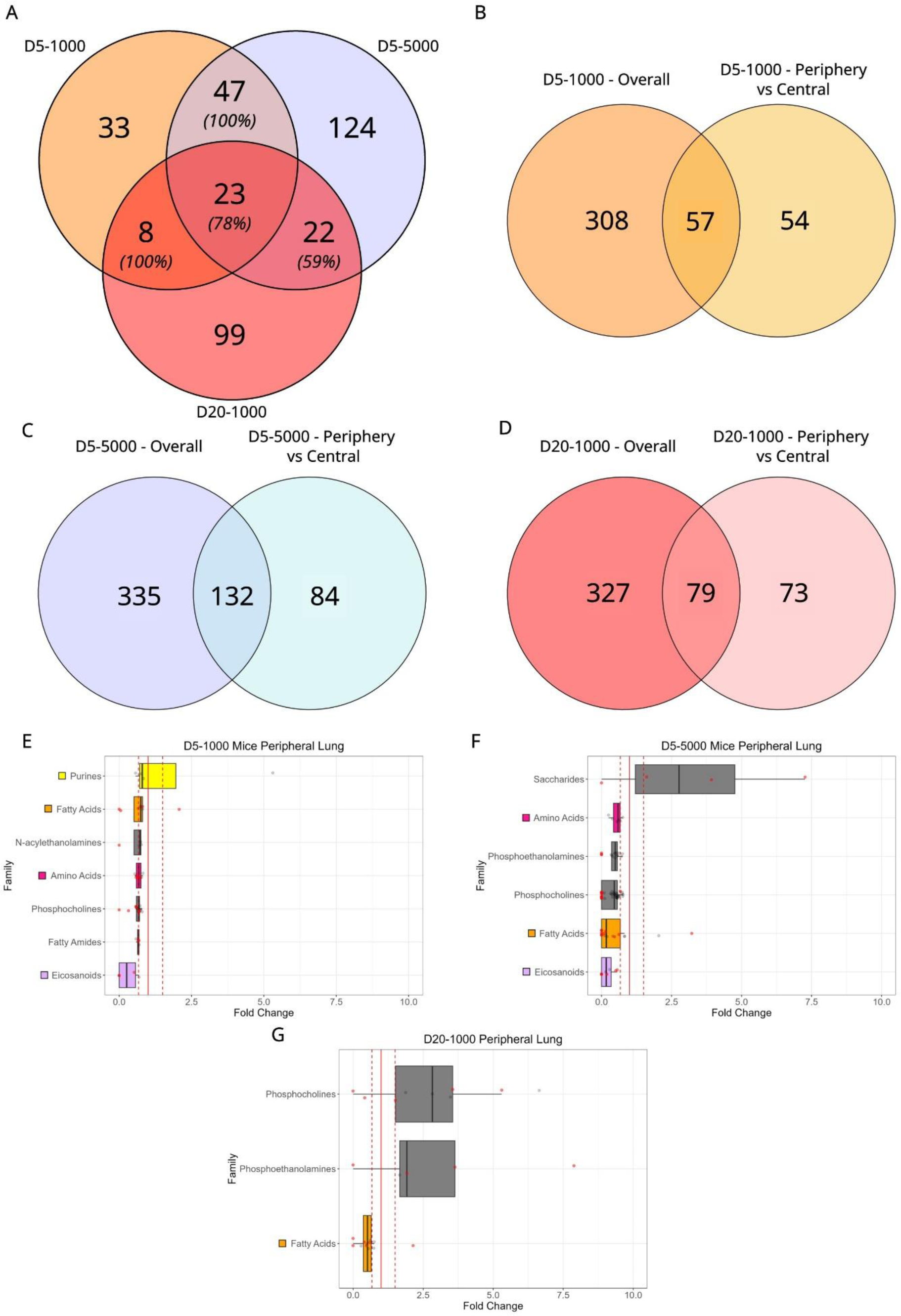
SARS-CoV-2 infection has differential localized metabolic effects on the metabolome of peripheral and central lung tissue. (A) Overlap of metabolites that are significantly different between peripheral and central lung tissue when normalized to their timepoint-matched mock-infected controls. Italicized percentages in each overlap are the percent of metabolites that all have the same direction of change. (B-D) Comparison of the metabolites that significantly differ between infected and uninfected mice without consideration of position and metabolites that are significantly different between peripheral and central lung tissue when normalized to their timepoint-matched mock-infected controls in (B) D5-1000, (C) D5-5000, and (D) D20-1000. (E-G) Significantly altered molecular families for the peripheral lung tissue of (E) D5-1000, (F) D5-5000, and (G) D20-1000 mice when normalized to their timepoint-matched mock-infected controls and when compared to central lung tissue. All families depicted had at least 4 annotated metabolites belonging to that family. Dots on the right edge of each panel represent metabolites that were only detected in infected samples (fold change was infinite) or had fold changes greater than 5. Dashed red lines represent a fold change of 0.67 or 1.5 while the solid red line represents a fold change of 1. Individual family members that were not significantly different overall but were significantly different between the peripheral and central tissue are colored red while metabolites that were significantly different overall in addition to being significantly different between peripheral and central tissue are gray. Family colors as in Figure 2. All data represented as median fold change.

Levels of tryptophan (Figure 5A), tyrosine, and phenylalanine were all significantly more decreased by infection in peripheral lung segments than central lung segments in D5-1000 and D5-5000 mice. Prostaglandin I_3_ (Figure 5B) and an unannotated arachidonic acid-derived prostaglandin (Figure 5C) were both significantly lower in the peripheral tissue of D5-1000 and D5-5000 mice. D5-5000 mice also had lower levels of PGF_2α_ and an unannotated eicosapentaenoic acid-derived prostaglandin in their peripheral tissue. Docosahexaenoic acid (Figure 5D) was significantly higher in peripheral tissue when compared to central tissue. The fatty amides palmitamide and octadecanamide (Figure 5E) were significantly lower in peripheral tissue of D5-1000 and D5-5000 mice when compared to central tissue. Lastly, a number of phosphocholines were at significantly lower levels in peripheral tissue when compared to central tissue in both D5-1000 and D5-5000 mice (Figure 4E-F; 5F). Fold differences and p-values for all mentioned metabolites can be found in Supplemental Table 1.

**Figure 5.**
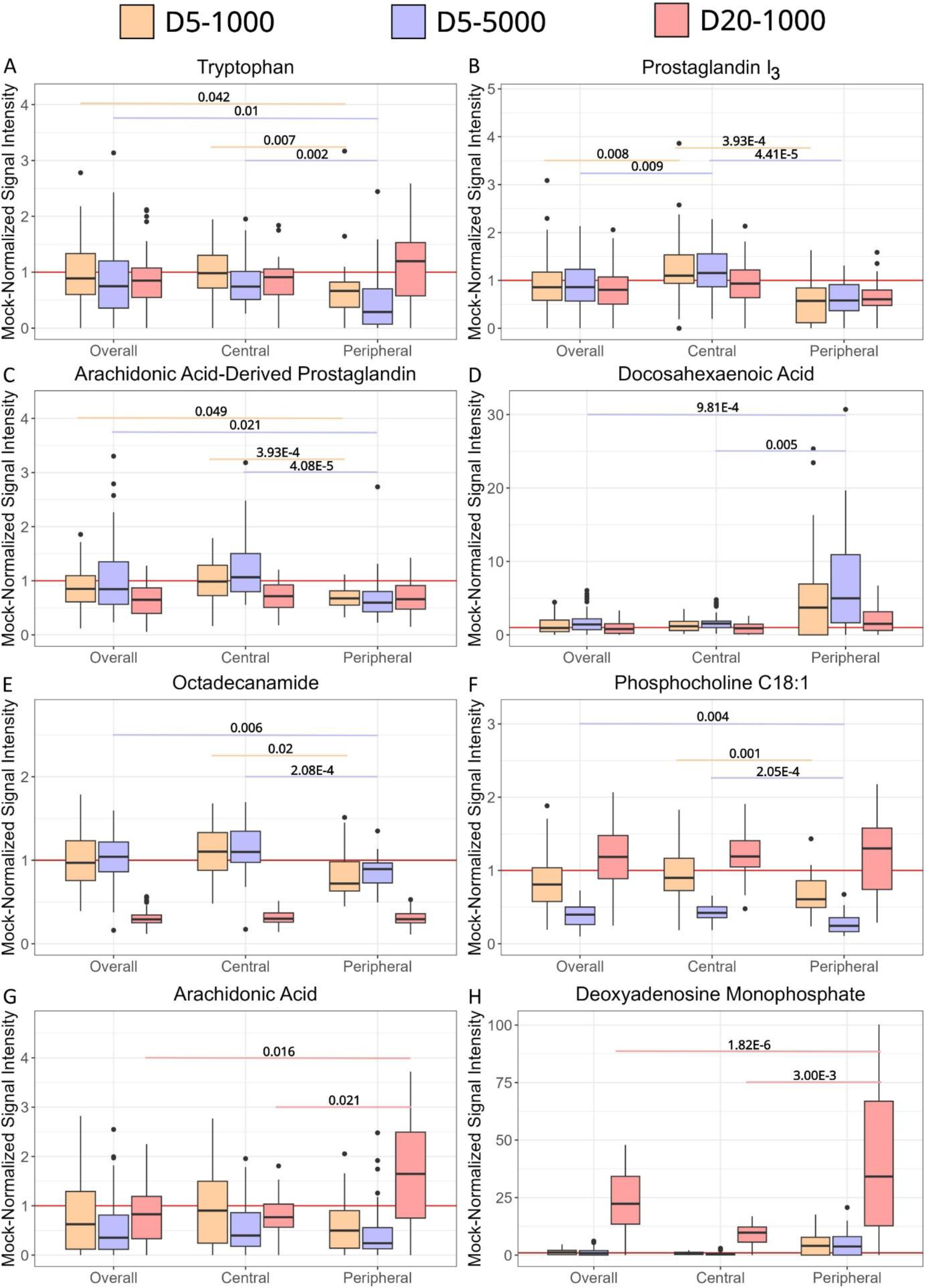
SARS-CoV-2 infection has localized effects on individual metabolites. (A-H) Boxplots comparing impact of infection on overall, central, and peripheral levels of (A) tryptophan, (B) prostaglandin I3, (C) an unannotated arachidonic acid-derived prostaglandin (*m/z* 353.23), (D) docosahexaenoic acid, (E) octadecanamide, (F) phosphocholine C18:1, (G) arachidonic acid, and (H) deoxyadenosine monophosphate in D5-1000 (orange), D5-5000 (purple), and D20-1000 mice (red). All metabolite signal intensities were normalized to the median signal intensity of the mock-infected mice for each position. P-values were calculated for intergroup comparisons for each disease state and were adjusted using the Bonferroni method. Only significant p-values are shown. The red line represents a fold difference of 1.

The impact of infection in D20-1000 mice demonstrated different spatial patterns than in D5-1000 and D5-5000 mice. Amino acids and eicosanoids exhibited comparable impact of infection in peripheral and central segments (Supplemental Table 1). Several fatty acids did not demonstrate any substantial localized effects in D5-1000 or D5-5000 mice; however, the impact of infection on arachidonic acid was two times higher in peripheral levels than central levels in D20-1000 mice (Figure 5G). Deoxyadenosine monophosphate (Figure 5H) was over 3 times higher in peripheral lung tissue than central lung tissue in D20-1000.

### Peripheral and central lung tissue shows innate differences in metabolite distribution prior to SARS-CoV-2 infection

Since SARS-CoV-2 virus demonstrated preferential localization to central lung tissue while infection was associated with higher levels of metabolic perturbance in the peripheral lung tissue, we sought to understand factors that could contribute to viral and metabolic localization. When analyzing the periphery of the mock-infected lungs, mice innately had higher levels of fatty amides while having lower levels of amino acids, acylcarnitines, phosphocholines, glycerolipids, and eicosanoids (Figure 6A). Fatty acids were also significantly altered but presented with mixed directions of change in the periphery.

**Figure 6.**
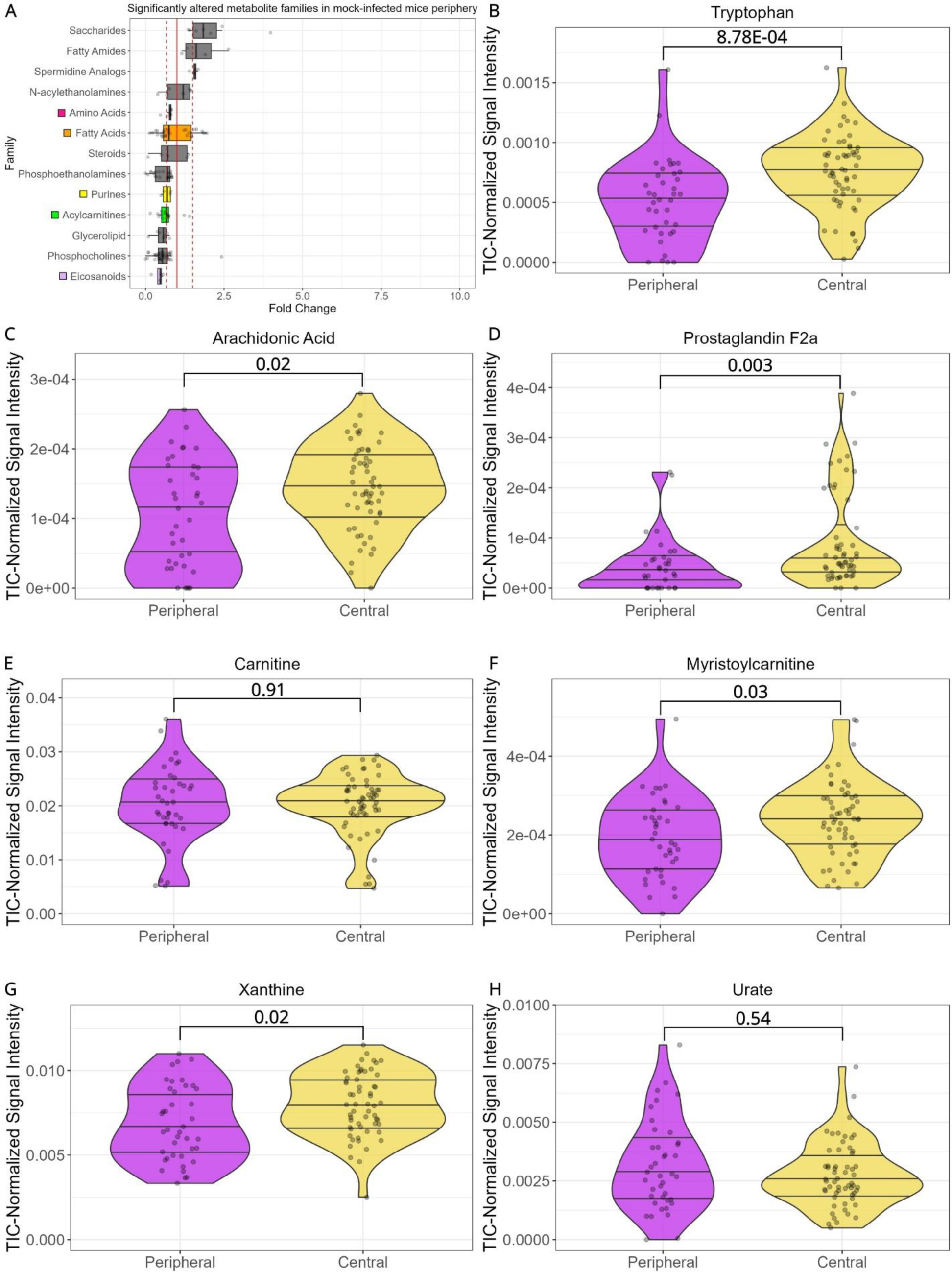
Peripheral and central lung tissue shows innate differences in metabolite distribution prior to SARS-CoV-2 infection. (A) Significantly altered molecular families between central mock-infected lung tissue and peripheral mock-infected lung tissue. All families depicted had at least 4 annotated metabolites belonging to that family. Dots on the right edge of each panel represent metabolites that were only detected in infected samples (fold change was infinite) or had fold changes greater than 5. Dashed red lines represent a fold change of 0.67 or 1.5 while the solid black line represents a fold change of 1. Family colors as in Figure 2. Data represented as median fold change (B-H) Violin plots of individual metabolites of interest.

Tryptophan (Figure 6B), tyrosine, phenylalanine, and methionine were all significantly lower in peripheral tissue when compared to central tissue (Supplemental Table 1). While both arachidonic acid (Figure 6C) and docosahexaenoic acid were significantly lower in the peripheral lung tissue, eicosapentaenoic acid was not significantly different between central and peripheral lung tissue. Of the previously discussed eicosanoids, only PGF_2α_ showed any spatial segregation, with central segments showing higher levels relative to peripheral segments (Figure 6D). With the exception of acetylcarnitine, acylcarnitines tended to be lower in peripheral tissue (Supplemental Table 1). Myristoylcarnitine (Figure 6F) and hexanoylcarnitine were both significantly lower while carnitine (Figure 6E) and oleoylcarnitine were not significantly different between the lung segments. Guanine, guanosine, and xanthine (Figure 6G) were all significantly lower in peripheral tissue, but urate (Figure 6H) did not show any significant differences.

## Discussion

Currently, metabolomics studies of COVID-19 have focused on the analysis of biofluids (plasma, serum, etc.), but biofluids cannot fully encapsulate the intricacies of metabolism in the lungs. Indeed, analysis of lung tissue in other infection systems has revealed new findings not observed through biofluids or that contradict those found in a biofluid.^40,41^ To address this gap, we systematically analyzed lung samples from a mouse model of SARS-CoV-2 infection, using mouse-adapted SARS2-N501Y_MA30_ virus.^28^ In this study, infection at two different viral doses and two different timepoints demonstrated a depleting effect on metabolism, with the majority of metabolites showing a decrease compared to mock-infected samples (Figure 1F-H). This stands in contrast to findings in plasma and serum, where acylcarnitines^21,42,43^, the amino acids phenylalanine and methionine^44–46^, and purines^47,48^ were all elevated with COVID-19.

COVID-19 unevenly impacts the lower respiratory system, particularly the lower lobes of the lungs.^13,14^ A strength of spatial metabolomics (“chemical cartography”) studies is the ability to define how different regions of a given tissue respond to infection.^49^ Implementation of this approach in this study demonstrated localized effects of SARS-CoV-2 infection in central and peripheral lung tissue, with the strongest metabolic impact observed in the lower left lung (Figure 3). Post-infection viral load tended to be highest in central lung tissue, while metabolic disturbance was highest in peripheral tissue. Fatty acids, amino acids, and eicosanoids were all more strongly decreased by infection in the peripheral D5-1000 and D5-5000 lung tissue than in central lung tissue (Figure 4). These localized effects of metabolism could help explain why peripheral lung tissue tends to be impacted more by COVID-19.

It is well-known that amino acids show signs of dysregulation in biofluids from patients with COVID-19 and other viral conditions.^50–52^ We observed this trend of amino acid dysregulation in the lungs overall (Figure 2A-D), with the majority of significantly altered amino acids showing decreases upon infection, but we also discovered spatial trends in amino acid distribution. Mock-infected mice exhibited higher levels of amino acids in the central portions of the lung when compared to the peripheral portion (Figure 6A-B). In infected mice, amino acids were more strongly decreased by infection in the peripheral lung when compared to the central lung (Figure 4E-G; 5A). Higher availability of amino acids in the central lung could make it easier for SARS-CoV-2 to replicate, leading to higher viral loads in these portions of the lung. Increases to the ratio of kynurenine to tryptophan have been associated with severity of COVID-19.^53,54^ The significant decrease in tryptophan and significant increase in kynurenine associated only with D5-5000 mice corresponds well with the 5000 pfu dosage being lethal and the 1000 pfu dosage being survivable (Figure 2D; Supplemental Figure 4). Glutamine, glutamate, and aspartate all significantly correlated with viral load across D5-1000 and D5-5000 mice combined (spearman rho = 0.34, 0.44 and 0.53, respectively). These three amino acids are all part of glutamine metabolism, which is increased in immune cells like macrophages and lymphocytes.^55,56^ The correlation between glutamine metabolites and regions rich with viral load could be a result of the recruitment of immune cells to fight off viral infection and proliferation.

Prostaglandin D_2_ and prostaglandin F_2α_ have a complicated role in immune response regulation across the body with a mixture of both pro-inflammatory and anti-inflammatory responses.^28,32,57–61^ PGD_2_ and PGF_2α_ were lower overall in D5-1000 mice and D20-1000 mice, but significantly higher in D5-5000 mice (Figure 2F); arachidonic acid, the precursor to PGD_2_ and PGF_2α_, was significantly lower in D5-5000 mice, which has been observed in other studies of COVID-19 (Figure 2E).^62^ In murine SARS-CoV-2 infection, PGD_2_ signaling had deleterious effects on animal survival, viral clearance, and lung edema.^28^ PGD_2_ increases in the lungs as a natural result of aging, amplifying the increase of PGD_2_ displayed with increased disease severity and potentially serving as one of the factors for increased mortality in elderly patients.^28^ On the other hand, prostaglandin I_3_, an anti-inflammatory eicosanoid and key mediator in inflammatory responses^63^, was not significantly altered overall. Despite this, PGI_3_ levels were significantly lower in D5-1000 and D5-5000 peripheral lung tissue when compared to central lung tissue (Figure 5B). While not directly related to anti-inflammatory eicosanoid production, docosahexaenoic acid (DHA) was significantly higher in peripheral tissue of D5-5000 mice (Figure 5D). DHA serves as a precursor for anti-inflammatory protectins and resolvins^64^, so higher levels of free DHA could be a result of decreased production of these anti-inflammatory products. A healthy inflammatory response requires a balanced mixture of ω-3 and ω-6 derived eicosanoids, protectins, and resolvins.^65^ In D5-5000, we see a clear preponderance of disease-promoting and pro-inflammatory metabolites in the overall lung without an equal response in anti-inflammatory metabolites. This imbalance appears to be worse in the peripheral lung tissue, with lower levels of anti-inflammatory molecules than central lung tissue, which could help explain the increased metabolic disturbance in this region. The use of ω-3 fatty acid supplements of DHA and EPA have been proposed as a method for combating severe COVID-19^64,66,67^, but double-blind clinical studies investigating this treatment method either do not have publicly available results or have been terminated due to difficulty with patient enrollment.^68–70^

Infected mice had significantly lower levels of purine nucleosides. Viruses are known to upregulate metabolites necessary for their replication such as nucleotides and amino acids. Here, we see a pattern of depletion on these groups, possibly indicating incorporation into viral macromolecules, leading to lower levels of free nucleotides and amino acids detectable by metabolomics analysis. In D5-5000 mice, the end product of purine metabolism, urate, was significantly increased. Cell death leads to increased production of urate, which in turn causes a pro-inflammatory response.^71–73^ Additionally, the production of urate from purines causes the release of radical oxidative species, which can cause oxidative-stress.^74^ Gout, an inflammatory joint condition marked by increased levels of urate, has been linked to increased likelihood of COVID-19 and COVID-19 disease severity.^75–77^ While this has been theorized to be a result of comorbidities, here we see that severe COVID-19 disease results in elevated urate levels in the lungs, which could compound upon existing hyperuricemia and lead to worse patient outcomes. Colchicine, an anti-inflammatory drug sometimes prescribed to gout patients, has shown some success in reducing severity of COVID-19 in patients with PCR-confirmed cases^78^, but urate lowering therapy has not shown to reduce COVID-19 severity.^76,79^ The greater efficacy of generalized anti-inflammatory treatments compared to treatments focused purely on urate reduction is consistent with the fact that the increased levels of urate are just one of several pro-inflammatory metabolite changes in SARS-CoV-2 infected mice.

D20-1000 mice still demonstrated significantly different metabolism from mock-infected mice and significantly higher viral loads, although to a lesser degree than D5-1000 mice (Supplemental Figure 1). Over half of the metabolites significantly altered in D5-1000 and D5-5000 were significantly altered in D20-1000, including key amino acids, fatty acids, and acylcarnitines (Figure 1C). The primary difference between D20-1000 and D5-1000 mice appeared to be in purine metabolism. Guanosine monophosphate was detected in D5-1000 and D5-5000 mice, but not in D20-1000 mice, indicating lowered levels. In addition to lower guanosine monophosphate, D20-1000 had significant increases in guanine and guanosine not present in D5-1000 mice, indicating decreased usage of the purine salvage pathway. This could be a result of decreased viral replication at this stage of infection, which would decrease the need for GMP/AMP and therefore increase free purines, or it could be an overcompensation of purines in response to the purine depletion seen in D5-1000 mice.

In spite of the strong genetic similarity of inbred C57BL/6J mice, D20-1000 mice presented with a range of recovery responses, with some mice showing overall metabolic recovery, and others showing persistently perturbed metabolism (Supplemental Figure 2A). Lung recovery also varied based on lung segment; upper lung segments of D20-1000 mice had similar metabolic profiles to upper lung segments of D5-1000 mice, while lower lung segments displayed significant metabolic differences that trended towards metabolic recovery, suggesting uneven healing in the lungs as the infection resolves (Supplemental Figure 2B; Supplemental Table 4). Jointly, these results may represent a path towards understanding the heterogeneity in long COVID development, though studies of later time points will be necessary. Future studies of COVID-19 recovery and long COVID should also investigate the metabolism of the brain, pancreas, and heart, based on symptoms seen during acute and post-acute COVID-19.

This study used the mouse-adapted COVID-19 strain SARS2-N501Y_MA30_ for infection in wild-type mice. While this strain of SARS-CoV-2 does provide a good experimental model for COVID-19 infection^28^, it is neither the same strain of SARS-CoV-2 that infects humans, nor the latest strain of SARS-CoV-2 since the virus mutates rapidly. Nevertheless, we observed many similarities between the chemical classes that are significantly altered in human studies and in our data.^80–82^ Additionally, the localized effects in peripheral lung tissue observed in this model match those seen in human findings.^14,16–18^ Differences between human data and this study may be due to the use of a mouse model, or to the fact that most human studies were restricted to biofluids, whereas our mouse model enabled the analysis of lung tissue, the primary site of infection for SARS-CoV-2. Future studies should investigate peripheral lung metabolism in the transverse plane, as this region frequently shows ground-glass opacities on x-ray and CT scans.^83^ To our knowledge, this is the first study that performed spatial metabolomics analysis on SARS-CoV-2 infected lungs. We observed higher viral loads in central lung tissue. In spite of this, metabolic changes were largest in peripheral lung tissue, matching the higher levels of peripheral lung tissue damage observed in humans. Mock-infected mice had innately higher levels of acylcarnitines, amino acids, fatty acids, and purines in their central lung tissue, which might aid in SARS-CoV-2’s localization to centralized lung tissue. Infected lung tissue presented with significantly lower levels of acylcarnitines, fatty acids, most eicosanoids, and amino acids. Several fatty acids and amino acids demonstrated significantly different levels of metabolites in peripheral lung tissue, where the largest degree of lung damage is observed in humans. These findings, which were only achieved due to spatial analysis of lung tissue, help contextualize localized effects of COVID-19 in the lungs and provide insight on potential molecular families to target for future mitigation of COVID-19 symptoms.

## Acknowledgements

This project was supported by start-up funds to L-I.M and by funds from the National Institutes of Health (P01 AI060699 and R01 AI129269) to SP. L-I.M. holds an Investigators in the Pathogenesis of Infectious Disease Award from the Burroughs Wellcome Fund.

## Author Contributions

Conceptualization, L-I.M.; Methodology, J.R., S.P. and L-I.M.; Software, J.R.; Formal Analysis, J.R.; Investigation, J.R., B.X., J.Z., and M.N.; Resources, S.P. and L-I.M.; Data Curation, J.R.; Writing – Original Draft, J.R.; Writing – Review & Editing, J.R., B.X., J.Z., M.N., S.P., and L-I.M.; Visualization, J.R.; Supervision, L-I.M.; Project Administration, L-I.M.; Funding Acquisition, S.P. and L-I.M.

## Declaration of Interests

The authors declare no competing interests.

## Supplemental information

Document S1. Figures S1–S6 and Tables S2 – S9

Excel S1. Excel file containing data for Table S1 which is too large to fit in a PDF

## Materials and Methods

### Ethics statement

All vertebrate animal studies were performed under IACUC protocol number 2071795, approved by the University of Iowa Institutional Animal Care and Use Committee.

### Viral strain

Mice exhibit resistance to ancestral SARS-CoV-2 infection. To allow for murine infection, a point mutation was introduced into SARS-CoV-2 (SARS2-N501Y_MA30_), as described in Wong et al.^28^ After 30 serial passages through mouse lungs, the modified virus demonstrated increased adaptation and lethality in mice.^28^ Plaque-purified clone from the modified virus was further propagated in Calu-3 2B4 cells and used to infect mice.

### Mouse infection

Nine-month-old female C57BL/6J mice were intranasally inoculated with 1000 PFU (n=12, sublethal dose) or 5000 PFU (n=6, lethal dose) of SARS2-N501Y_MA30_ virus in a total volume of 50 μl DMEM. 6 mice infected with the sublethal dose and 6 mice infected with the lethal dose were anesthetized with ketamine-xylazine at day 5 post-infection for lung collection. 6 mice infected with the sublethal dose were anesthetized at day 20 post-infection. 5 mock-infected mice were sacrificed at day 5 and day 20 respectively as time-matched mock controls. Mouse lungs were harvested and divided into 12 pieces as shown in Figure 3B for metabolic analysis and viral load analysis.

### Metabolite Extraction

Samples were snap frozen, weighed and homogenized in LC-MS grade water (10 µL/mg) with steel beads using a Fisherbrand Bead Mill 24 Homogenizer at a speed of 5 m/s for 20 seconds. 1/10 of the homogenized samples was saved for RNA extraction, while the rest of the volume was extracted for metabolites according to Want et al.^84^ Briefly, LC-MS grade methanol was added to each sample for a final concentration of 50%. Samples were homogenized again for 20 seconds and centrifuged at 16,000g for 10 minutes at 4℃. The 120 µL of the supernatant (aqueous extract) was collected, subjected to overnight vacuum drying using a Speedvac, and then frozen at −80℃ until LC-MS analysis. A mixture of dichloromethane and methanol at a ratio of 3:1 (V/V) was added to the aqueous extraction pellet. Samples were homogenized for 20 seconds, centrifuged at 16,000g for 10 minutes at 4℃, dried overnight (120 µL) and frozen at - 80℃ until LC-MS analysis.

### Viral load analysis

To quantify the titers of SARS-CoV-2 virus in the indicated sections of lungs from mock-infected and infected mice, 1/10 volume of homogenized samples was collected for RNA extraction by using TRIzol Reagent. 0.2 μg total RNAs were converted to cDNA using High-Capacity cDNA Reverse Transcription Kit (ThermoFisher, cat#: 4368814). RT-qPCR was conducted by using SYBR Green real-time PCR master mix (ThermoFisher, cat#: A25778) in 20 μl reactions using standard reaction conditions (50℃/2min, 95℃/10min, 95℃/15sec, 60℃/1min) with 40 PCR cycles. Average values from duplicates of each gene were used to calculate the relative abundance of transcripts normalized to mouse GAPDH and presented as 2^−ΔΔCT^. The primers used for amplification of viral N gene were reported previously.^28^

### Metabolite resuspension and LC-MS/MS data collection

Both the organic and aqueous extracts were resuspended in 75 µL of 50:50 LC-MS grade methanol:water spiked with 2 µM sulfadimethoxine. Samples were sonicated for 5 minutes at a frequency of 40 kHz using a Fisher Scientific ultra-sonicator. Samples were then centrifuged for 5 minutes at 12.8G. 60 µL of the supernatant from the organic and aqueous portions of each sample were combined into a single well of a 96-well plate, resulting in 120 µL volume prior to analysis. Extraction blanks were resuspended in the same way. A pooled quality control (QC) was created by combining 2 µL of each sample. A randomized run sequence was generated through the use of the RAND() function in excel to minimize batch effects. LC analysis was performed using a Kinetex 1.7 µm C8 100 Å, 50 x 2.1 mm column (S/N H17-371701, B/V 5606-0131) on a Vanquish HPLC (Thermo Scientific). The column temperature was set to 40 °C. Mobile phase A was LC-MS grade water (Fisher Optima) with 0.1% formic acid (Fisher Optima, CAS 64-18-6) and mobile phase B was LC-MS grade acetonitrile (Fisher Optima) with 0.1% formic acid. Liquid chromatography gradient parameters, as per ^10^, can be found in Supplemental Table 6. MS/MS analysis was performed using a Q-Exactive Plus (Thermo Scientific) high resolution mass spectrometer with the parameters found in Supplemental Table 7, as per ^10^. Prior to MS/MS analysis, the instrument was calibrated using Pierce LTQ Velos ESI positive ion calibration solution (Thermo Scientific). A 2, 5, 10, 20, and 30 µL volume of the QC was injected to determine the ideal injection volume for these samples^85^. By examining the linearity of the signal intensity and peak shape of internal standards and metabolites of interest as well as the overall signal intensity of the sample, a sample injection volume of 5 µL was determined to be the optimal injection volume. 5 µL of a blank and QC were injected between every 12 samples to monitor batch effects and signal drift. A mixture of 6 internal standards were injected at the start of the run, approximately every 100 samples, and at the conclusion of the run to monitor column health. No signs of batch effects or column deterioration were observed during the course of the run.

### Raw Data Processing

Following data collection, raw spectra were converted into the mzML format using MSconvert ^86^ before being analyzed using MZmine 2.53.^87^ Parameters for MZmine were adjusted by manually inspecting built chromatograms within the samples for signs of over and under splitting. A full list of parameters can be found in Supplemental Table 8. Metabolites that lacked an average signal intensity greater than 3 times the intensity of the metabolite in blanks were removed. Signal intensity was then normalized using the Total Ion Current (TIC) normalization method.

### Data Analysis

QIIME 2^88^ was used to perform all calculations related to the Principal Coordinate Analysis (PCoA). Distance matrices were calculated using “diversity beta” with the Bray–Curtis dissimilarity method on the TIC-normalized MS1 data. PCoAs were generated with “diversity pcoa” and visualized with “emperor plot”. Pseudo-F values were calculated based on the Bray-Curtis distance matrices at each position using “beta diversity group significance”.

All visualizations of the lung were generated using ‘ili ^89^, on the lung model generated in _10_.

Feature-based molecular networking was performed using the global natural products social molecular networking resource (GNPS).^90,91^ A full list of parameters used to perform FBMN can be found in Supplemental Table 9, as per ^10^. Level 2/3 metabolite annotations (as outlined in the Metabolomics Standards Initiative rankings^92^) were achieved through the use of FBMN and manual inspection of mirror plots to ensure a quality match of MS2 spectra. Level 1 metabolites annotations were assigned by matching retention times of each annotation with LC-MS grade analytical standards using the same column and LC gradient. A full list of metabolite annotation confidence levels for each metabolite mentioned can be found in Supplemental Table 1.

Chemical families were assigned using annotations and FBMN. Canonical SMILES for annotated metabolites were identified using NPClassifier.^93^ A general family assignment was given to all members of each FBMN family based on the annotated metabolites in each FBMN family. For annotated metabolites with statistical significance, families were manually validated.

### Statistical Analysis

All statistical analysis was performed using R version 4.1. To determine statistical significance between two groups, groups were first tested for normality using the shapiro.test() function in version 4.1.2 of the “stats” package in R. Groups that both had a normal distribution were tested for significance using the t.test() function, while non-normally distributed groups were tested with wilcox.test(). Correlation analysis was performed using the Spearman method in the cor.test() function in version 4.1.2 of the “stats” package in R. False-discovery-rate adjusted p-values were generated from the entire list of p-values in each analysis using the p.adjust() function with the “fdr” method in version 4.1.2 of the “stats” package in R. Violin plots and boxplots were generated using version 3.4.3 of “ggplot2”. χ^2^ values were generated using the chisq.test() function in base r. For chi-squared calculations, the expected distribution for upregulated versus downregulated was 0.5 and 0.5.

For comparisons between central and peripheral lung tissue, the TIC-normalized signal intensity for each metabolite in the infected tissue was divided by median value of the TIC-normalized signal intensity of the metabolite in the same region of the time-matched mock-infected control.

## Data availability

Raw data files have been deposited at MassIVE at MSV000094591. The final GNPS feature-based molecular network job can be accessed through the following URL: https://gnps.ucsd.edu/ProteoSAFe/status.jsp?task=0950a8281f6748e783e419ae5bdf6d85

## Code availability

A R markdown notebook of code used for the project can be found at https://github.com/jarrodRoachChem/covid19

## Notes

https://gnps.ucsd.edu/ProteoSAFe/status.jsp?task=0950a8281f6748e783e419ae5bdf6d85

https://massive.ucsd.edu/ProteoSAFe/dataset.jsp?task=0934d537663042a9bd63d513c1c28fe3

### Competing Interest Statement

The authors have declared no competing interest.

