## Supplemental figures and tables 2-9 for "SARS-CoV-2 infection unevenly impacts metabolism in the coronal periphery of the lungs"

### Supplemental information

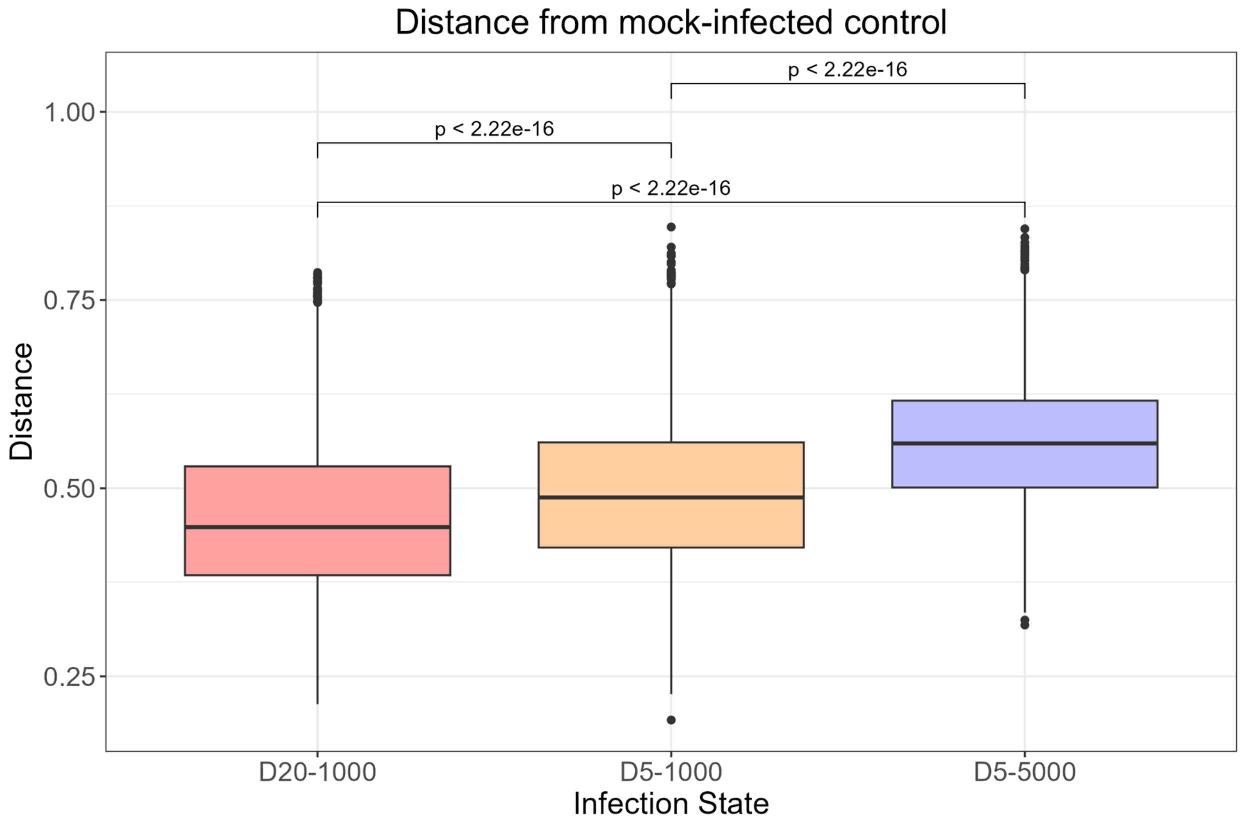


Supplemental Figure 1. Bray-Curtis metabolic distances between each disease state and its matched control.

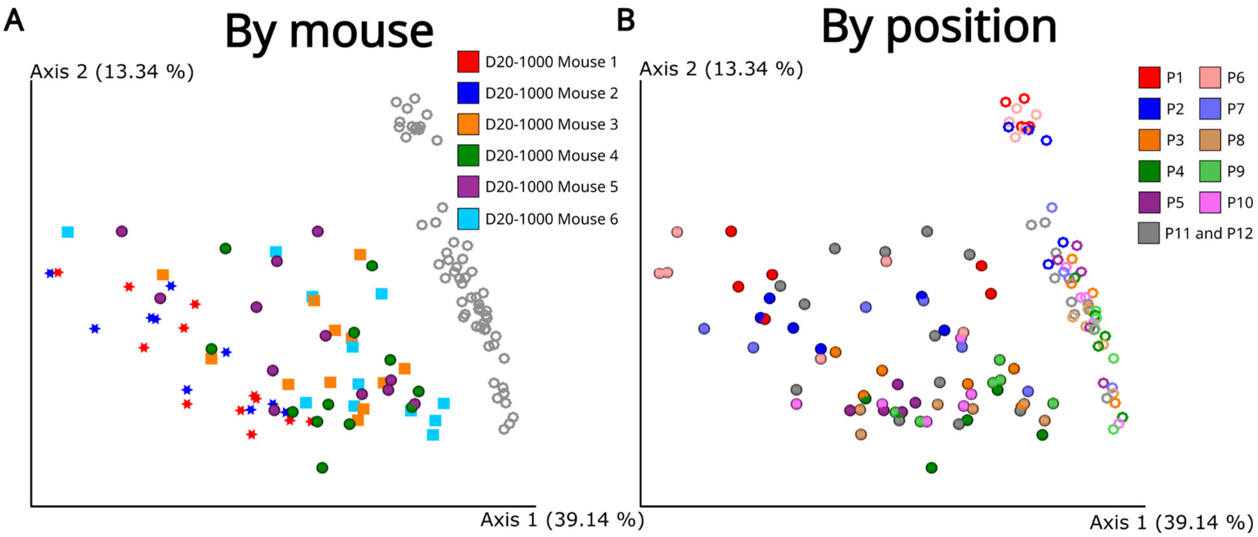


Supplemental Figure 2. D20-1000 mice and lung segments demonstrate a wide range of phenotypes. (A) Principal coordinate analysis with only D20-1000 (colored) and Mock20 (gray rings) visualized. Each color represents an individual D20-1000 mouse, with one sample for each of the 12 lung segments. Some mice (light blue and orange squares) trend more towards the mock-infected mice while others remain distant (dark blue and red stars). (B) Principal coordinate analysis with only D20-1000 (spheres) and Mock20 (rings) visualized. At D20, lower lung segments (P4, P5, P9, P10) trend towards mock-infected more than upper lung segments (P1, P2, P6, P7).

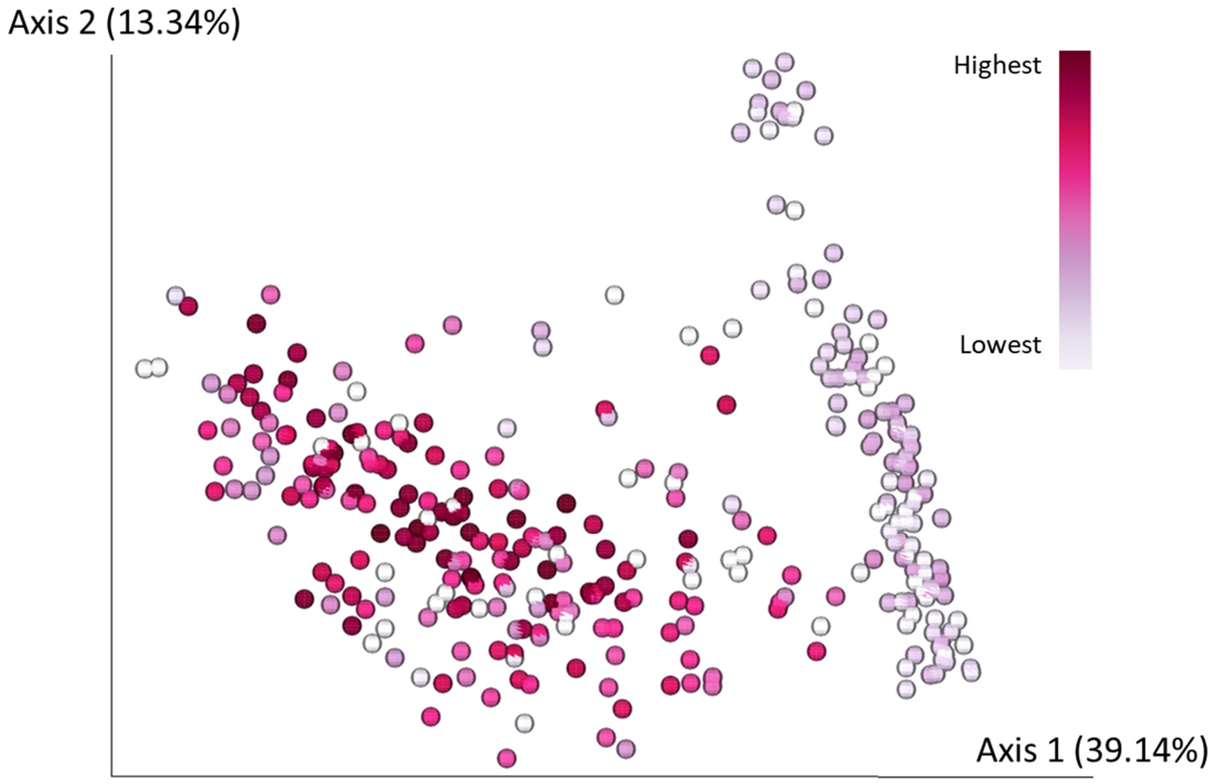


Supplemental Figure 3. Local tissue viral load does not correlate with overall metabolic effects of COVID-19.

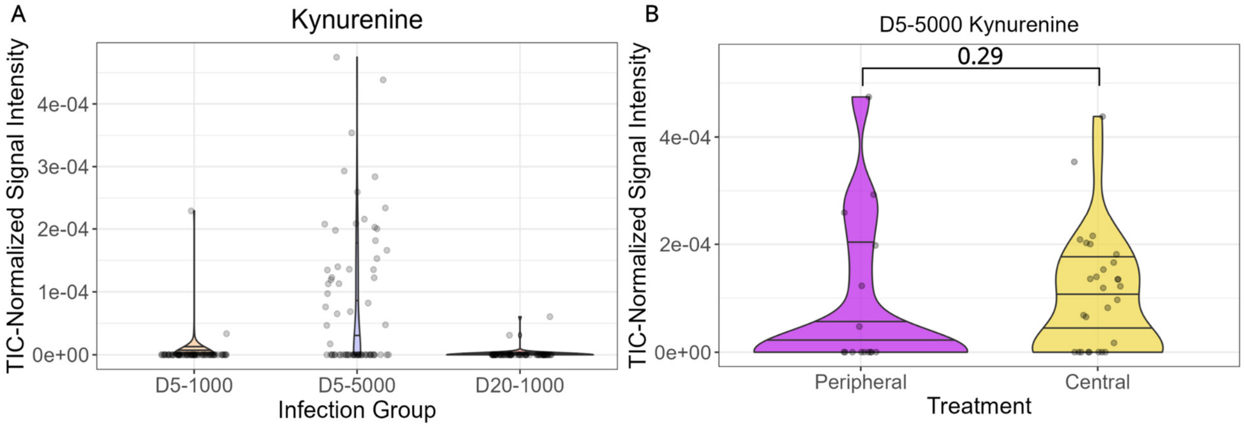


Supplemental Figure 4. Effects of SARS-CoV-2 on kynurenine. (A) TIC-normalized signal intensities for kynurenine for each lung segment in each infection state. Signal has not been normalized to mock-infected values as the signal intensity was 0 for all mock-infected lung segments. (B) Violin plot of kynurenine in peripheral and central lung segments of D5-5000 mice.

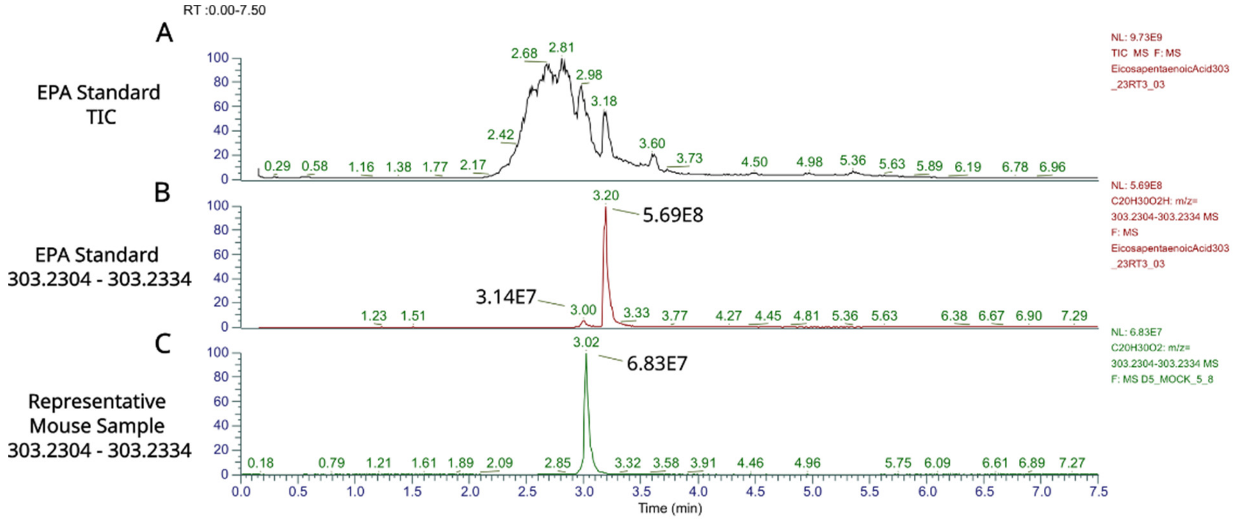


Supplemental Figure 5. Extracted ion chromatogram of eicosapentaenoic acid standard. (A) Total ion current for eicosapentaenoic acid standard. (B) Extracted ion chromatogram for eicosapentaenoic acid (monoisotopic *m/z* = 303.2324) from the EPA standard (C) Extracted ion chromatogram for eicosapentaenoic acid from a representative mouse sample.

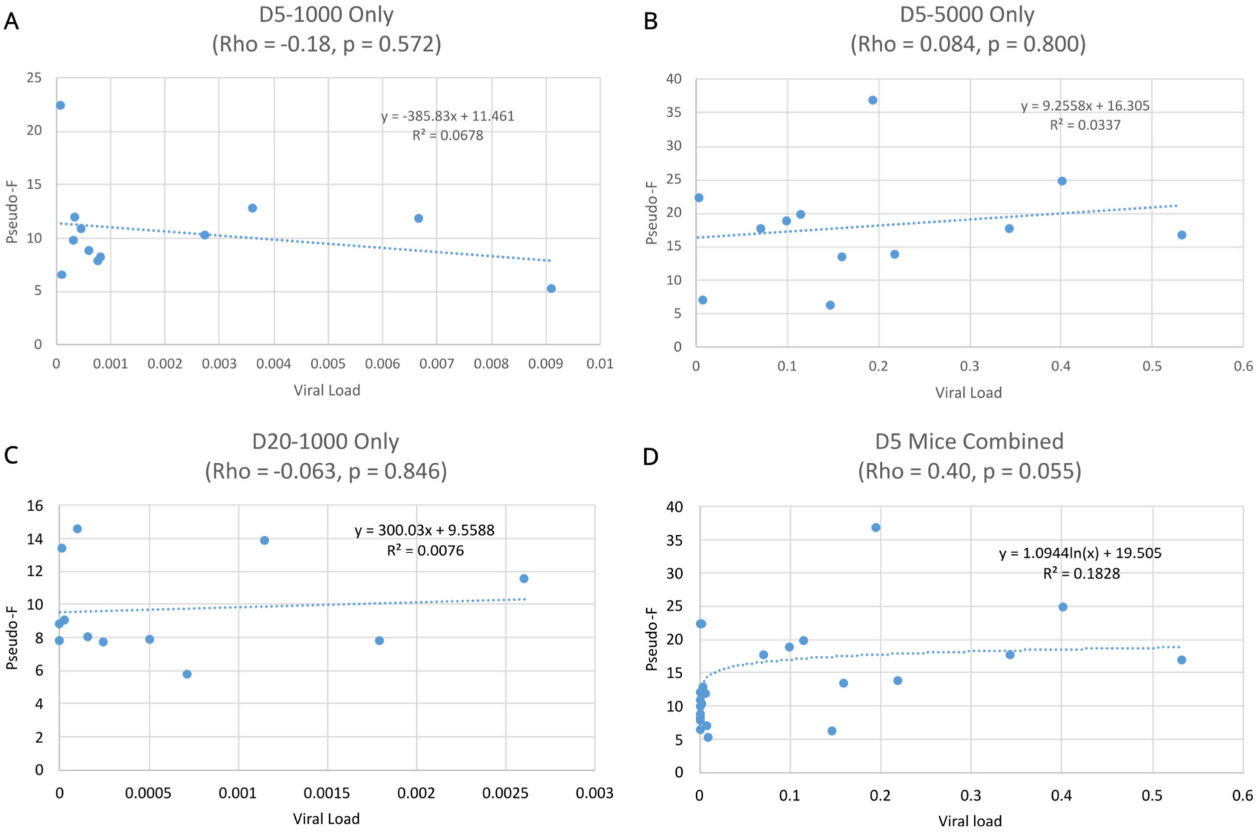


Supplemental Figure 6. Viral load does not correlate with pseudo-F values. Each panel shows the median viral load and the pseudo-F value associated with each lung segment (A) Only lung segments from D5-1000 mice. (B) Only lung segments from D5-5000 mice. (C) Only lung segments from D20-1000 mice. (D) Lung segments from D5-1000 and D5-5000 mice.

Supplemental Table 1. Fold change and FDR-adjusted p-values for metabolites mentioned by name (Excel file).

Supplemental Table 2. D5-5000 mice show the largest difference between peripheral and central lung segments. Q-values ≥ 0.05 are not considered significantly different.

| Experimental Group | Pseudo-F | Q-value |
|---|---|---|
| Mock5 | 3.6 | 0.001167 |
| D20-1000 | 5.0 | 0.005303 |
| Mock20 | 6.9 | 0.003316 |
| D5-1000 | 7.5 | 0.001167 |
| D5-5000 | 10.0 | 0.001167 |

Supplemental Table 3. Median viral loads, median distances between infected and time-matched mock-infected controls, and pseudo-F values.

| State | Position | Median Viral Load | Pseudo-F | Median Distance |
|---|---|---|---|---|
| D5-1000 | P1 | 0.003616527 | 12.87874 | 0.492648404 |
| D5-1000 | P2 | 0.002722801 | 10.36134 | 0.44057214 |
| D5-1000 | P3 | 0.006654237 | 11.91699 | 0.438300933 |
| D5-1000 | P4 | 0.00911257 | 5.297552 | 0.43209767 |
| D5-1000 | P5 | 8.42E-05 | 22.46072 | 0.482333338 |
| D5-1000 | P6 | 0.000588465 | 8.904733 | 0.5169872 |
| D5-1000 | P7 | 0.000336683 | 12.09997 | 0.46577933 |
| D5-1000 | P8 | 0.000318774 | 9.948068 | 0.344922064 |
| D5-1000 | P9 | 0.000818882 | 8.327394 | 0.389847885 |
| D5-1000 | P10 | 0.000451181 | 10.91874 | 0.477425786 |
| D5-1000 | P11 | 0.000754222 | 7.915017 | 0.487826178 |
| D5-1000 | P12 | 9.98E-05 | 6.641957 | 0.55254418 |
| D5-5000 | P1 | 0.114756468 | 19.99133 | 0.565713929 |
| D5-5000 | P2 | 0.09873198 | 18.95851 | 0.562750662 |
| D5-5000 | P3 | 0.400597292 | 24.93229 | 0.547479689 |

| | | | | |
|---|---|---|---|---|
| D5-5000 | P4 | 0.146384788 | 6.293912 | 0.453121262 |
| D5-5000 | P5 | 0.193794969 | 36.89948 | 0.597603795 |
| D5-5000 | P6 | 0.002329373 | 22.46339 | 0.614261757 |
| D5-5000 | P7 | 0.070131196 | 17.81124 | 0.547336842 |
| D5-5000 | P8 | 0.531234593 | 16.98055 | 0.472745007 |
| D5-5000 | P9 | 0.342997733 | 17.84118 | 0.469153544 |
| D5-5000 | P10 | 0.217984988 | 13.97498 | 0.530456642 |
| D5-5000 | P11 | 0.159078076 | 13.59289 | 0.516196791 |
| D5-5000 | P12 | 0.007401028 | 7.075114 | 0.488288134 |
| D20-1000 | P1 | 0.00260159 | 11.58302 | 0.475622558 |
| D20-1000 | P2 | 0.001147068 | 13.91599 | 0.48169412 |
| D20-1000 | P3 | 0.001789884 | 7.870275 | 0.400044511 |
| D20-1000 | P4 | 0.000152888 | 8.12082 | 0.389150647 |
| D20-1000 | P5 | 0.00010045 | 14.61029 | 0.414320013 |
| D20-1000 | P6 | 1.34E-05 | 13.48302 | 0.572041578 |

| | | | | |
|---|---|---|---|---|
| D20-1000 | P7 | 0.000715387 | 5.864108 | 0.407568469 |
| D20-1000 | P8 | 0.000506671 | 7.899302 | 0.36175649 |
| D20-1000 | P9 | 0.000242692 | 7.745196 | 0.332701188 |
| D20-1000 | P10 | 2.68E-05 | 9.083933 | 0.38287985 |
| D20-1000 | P11 | 0 | 7.861192 | 0.381681855 |
| D20-1000 | P12 | 0 | 8.857764 | 0.372799916 |

Supplemental Table 4. D20-1000 upper lung segments do not significantly differ from D5-1000 upper lung segments. Q-values ≥ 0.05 are not considered significantly different.

| Position | Lung Location | Pseudo-F | Q-value |
|---|---|---|---|
| P1 | Upper lung | 1.4 | 0.30 |
| P2 | Upper lung | 2.1 | 0.07 |
| P3 | Central lung | 2.7 | 0.03 |
| P4 | Lower lung | 3.9 | 0.02 |
| P5 | Lower lung | 4.8 | 0.02 |
| P6 | Upper lung | 1.3 | 0.25 |
| P7 | Upper lung | 1.4 | 0.24 |
| P8 | Central lung | 2.1 | 0.06 |
| P9 | Lower lung | 3.3 | 0.03 |
| P10 | Lower lung | 2.8 | 0.03 |
| P11 | N/A | 2.5 | 0.03 |
| P12 | N/A | 4.3 | 0.03 |

Supplemental Table 5. Lung segments in the same relative position on the left and right lung show similar overall metabolic profile. Q-values ≥ 0.05 are not considered significantly different.

| | **D5-1000** | | **D5-5000** | | **D20-1000** | | **Mock5** | | **Mock20** | |
|---|---|---|---|---|---|---|---|---|---|---|
| | **Pseudo-F** | **Q-value** | **Pseudo-F** | **Q-value** | **Pseudo-F** | **Q-value** | **Pseudo-F** | **Q-value** | **Pseudo-F** | **Q-value** |
| P1 and P6 | 0.9 | 0.47 | 1.4 | 0.25 | 1.0 | 0.38 | 0.9 | 0.47 | 0.8 | 0.74 |
| P2 and P7 | 1.2 | 0.3 | 0.9 | 0.59 | 0.4 | 0.8 | 0.6 | 0.88 | 3.3 | 0.05 |
| P3 and P8 | 0.5 | 0.82 | 1.5 | 0.17 | 0.8 | 0.52 | 1.9 | 0.1 | 0.5 | 0.75 |
| P4 and P9 | 1.1 | 0.39 | 0.2 | 0.98 | 0.7 | 0.71 | 0.8 | 0.77 | 0.6 | 0.6 |
| P5 and P10 | 1.5 | 0.2 | 2.3 | 0.06 | 0.7 | 0.67 | 1.3 | 0.29 | 0.6 | 0.59 |

Supplemental Table 6: Liquid Chromatography Gradient.

| Time | Flow [mL/min] | %B | Curve |
|---|---|---|---|
| 0.000 | 0.500 | 2.0 | 5 |
| 1.000 | 0.500 | 2.0 | 5 |
| 2.500 | 0.500 | 98.0 | 5 |
| 4.500 | 0.500 | 98.0 | 5 |
| 5.500 | 0.500 | 2.0 | 5 |
| 7.500 | 0.500 | 2.0 | 5 |

Supplemental Table 7: Q Exactive Plus (Thermo Scientific) instrument parameters.

| | |
|---|---|
| Run time | 0 to 7.5 min |
| Polarity | Positive |
| Default charge state | 1 |
| **Full MS** | |
| Resolution | 70,000 |
| AGC target | 3e6 |
| Maximum IT | 246 ms |
| Scan range | 100 to 1500 *m/z* |
| **dd-MS2** | |
| Resolution | 17,500 |
| AGC target | 1e5 |
| Maximum IT | 54 ms |
| Loop count | 5 |
| TopN | 5 |
| Isolation window | 1.0 *m/z* |

| | |
|---|---|
| (N)ce/stepped (N)CE | NCE: 20, 40, 60 |
| **dd Settings** | |
| Minimum AGC | 8e3 |
| Intensity Threshold | 1.5e5 |
| Peptide match | Preferred |
| Exclude isotopes | on |
| Dynamic exclusion | 10.0 s |

Supplemental Table 8: MZmine parameters.

| Type | Parameter | Value |
|---|---|---|
| Raw data methods > Feature detection > Mass detection | Mass detector | Centroid |
| | MS1 noise level | 4e5 |
| | MS2 noise level | 1e3 |
| Raw data methods > Feature detection > Chromatogram builder | Mass List | Previous Step Output |
| | Time span | 0.01 min |
| | Min height | 1.2e6 |
| | *m/z* tolerance | 10 ppm |
| Feature list methods > Feature detection > Chromatogram deconvolution | Local minimum | |
| | Chromatographic Threshold | 70% |
| | Search Minimum in Retention Time Range (minutes) | 0.1 |
| | Minimum Relative Height | 40.0 |
| | Minimum Absolute Height | 4.0E5 |
| | Minimum Ratio of Peak Top/Edge | 1.3 |
| | Peak Duration Range | 0.04 - 2.00 |
| | *m/z* MS2 | 0.01 |
| | RT MS2 (minutes) | 0.1 |
| Feature list methods > | *m/z* tolerance | 10 ppm |

| | | |
|---|---|---|
| Isotopes > Isotopic peak grouper | Retention Time Tolerance (minutes) | 0.1 |
| | Charge | 3 |
| | Representative isotope | Most intense |
| | Monotonic shape | Checked |
| Feature list methods > Alignment > Join aligner | m/z tolerance | 10 ppm |
| | Retention Time Tolerance (minutes) | 0.5 |
| | Weight *m/z* | 1 |
| | Weight RT | 1 |
| Feature List Methods > Filtering > Feature list rows filter | Minimum peaks in a row | 5 |
| | Retention Time | 0.2-7.50 |
| | Reset ID | Checked |

Supplemental Table 9: GNPS parameters.

| Feature-Based Molecular Network Parameters | | |
|---|---|---|
| Basic Options | Precursor Ion Mass Tolerance | 0.02 Da |
| | Fragment Ion Mass Tolerance | 0.02 Da |
| Advance Network Options | Min Pairs Cosine | 0.7 |
| | Minimum Matched Fragment Ions | 4 |
| | Maximum shift between precursors | 500 |
| | Network TopK | 10 |
| | Maximum Connected Component Size (Beta) | 100 |
| Advance Library Search Options | Library Search Minimum Matched Peaks | 4 |
| | Search analogs | Do Search |
| | Top results to report per query | 1 |
| | Score Threshold | 0.7 |
| | Maximum analog difference | 100.00 |
| Advanced Filtering Options | Minimum Peak Intensity | 0.0 |
| | Filter Precursor Window | Filter |
| | Filter peaks in 50 Da Window | Filter |

| | Filter Library | Filter Library |
|---|---|---|
| Advanced Quantification Options | Normalization Per File | Row Sum Normalization (Per file Sum to 1,000,000) |
| | Aggregation Method For peak abundances per group | Mean |
| Advanced External Tools | Run Dereplicator | Run |
